# Imprinted regulatory networks reveal the molecular crosstalk between paternal and maternal genomes in the endosperm of *Arabidopsis arenosa*

**DOI:** 10.64898/2026.07.23.740329

**Authors:** Trezalka Budinsky, Martin Kovačik, Vojtěch Čermák, Adéla Přibylová, Susnata Salony, Ömer Iltas, Aleš Pečinka, Clément Lafon Placette

## Abstract

Imprinted genes do not act alone to shape seed development, but as a complex network — just like any other gene. Yet, the molecular context in which they are embedded, i.e. their gene network, remains largely understudied. To address this knowledge gap, we characterized the imprintome of *Arabidopsis arenosa* at the species-level and used gene regulatory network analyses. We show that genomic imprinting preferentially affects only a few pathways, offering candidates for dosage sensitive processes and the molecular arena of parental conflict. In these pathways, some imprinted genes act as hub genes, among which *NRPE1*, highlighting the importance of epigenetics in endosperm development. The interaction between parental genomes was rather one-sided: paternally expressed regulators preferentially targeted PEGs, while maternally expressed regulators targeted both PEGs and MEGs indiscriminately. This aligns with a self-promoting paternal influence and a maternal buffer under a parental conflict scenario. Shared paternal and maternal regulation of downstream targets was nevertheless common, and we reveal novel molecular interactions between imprinted regulators. Overall, our work shows *how* genomic imprinting may act as a molecular means for parental conflict, but also highlights the importance of parental molecular coordination in endosperm development.

## Main

Genomic imprinting is an epigenetic mode of inheritance where alleles are unequally expressed based on their parent-of-origin^1^. It gives rise to maternally and paternally expressed genes (MEGs and PEGs), for which the paternal or maternal allele, respectively, is partially or completely silenced. The most prominent explanation for the evolution of genomic imprinting is the kinship (or parental conflict) model, which assumes that maternal and paternal evolutionary interests collide over the provisioning of the progeny. Consequently, it posits that MEGs and PEGs coevolve in a sexual arms race over resource allocation, with PEGs promoting resource allocation to the progeny at the expense of their half-siblings, and MEGs limiting resource allocation to the progeny to ensure fair resource distribution^2^.

In plants, a parallel “sexual key-lock” model assumes that the endosperm, which only forms upon fertilization, requires both maternal and paternal genomes for sustained development, with PEGs providing the components required to initiate endosperm development^3,4^. This evolutionary innovation in flowering plants would thus prevent wasteful resource allocation to an unfertilized ovule, unlike gymnosperms, where nourishing tissue arises prior to fertilization^5,6^. In the sexual key-lock model, MEGs and PEGs would work in synergy to facilitate endosperm development like two puzzle pieces coming together. The primary difference with the kinship model thus lies in the selective forces driving the coevolution of MEGs and PEGs. In the kinship model, MEGs and PEGs are fundamentally in conflict and genomic imprinting likely affects small-effect hence evolvable genes, while in the sexual key-lock model, their relationship is fundamentally cooperative, and imprinted genes would rather have pleiotropic effects.

Much of our current knowledge of genomic imprinting has been built from detailed studies of individual imprinted genes^5,7–9^. These studies have provided invaluable functional insights but have also produced an increasingly fragmented picture of genomic imprinting. We understand the functions of several prominent imprinted loci, yet we know comparatively little about how imprinted genes interact within the broader transcriptional landscape of the developing endosperm. This represents an important conceptual gap since gene products rarely function independently but rather operate within interconnected regulatory networks. In addition, focusing on a few genes can only give a limited view on the evolutionary drivers acting on genomic imprinting.

In this study, we characterized the imprintome of the diploid species *Arabidopsis arenosa*, a close outcrossing relative to the well-studied *A. thaliana*. We built gene regulatory networks (GRNs) to decipher the molecular context in which imprinted genes act. We characterized 30 imprintomes from 19 populations spanning the majority of the distribution and main phylogenetic lineages of diploid *A. arenosa*^10^ (Fig. 1a; Supplementary Table 1) and identified 39 MEGs and 45 PEGs found in all three phylogenetic lineages (Supplementary Table 2; Methods). We then investigated these imprinted genes in the context of their GRNs. Using weighted gene co-expression network analysis (WGCNA^11^), we constructed a GRN based on the same endosperm transcriptomes we used to identify imprinted genes (n = 62 samples).

**Fig. 1.**
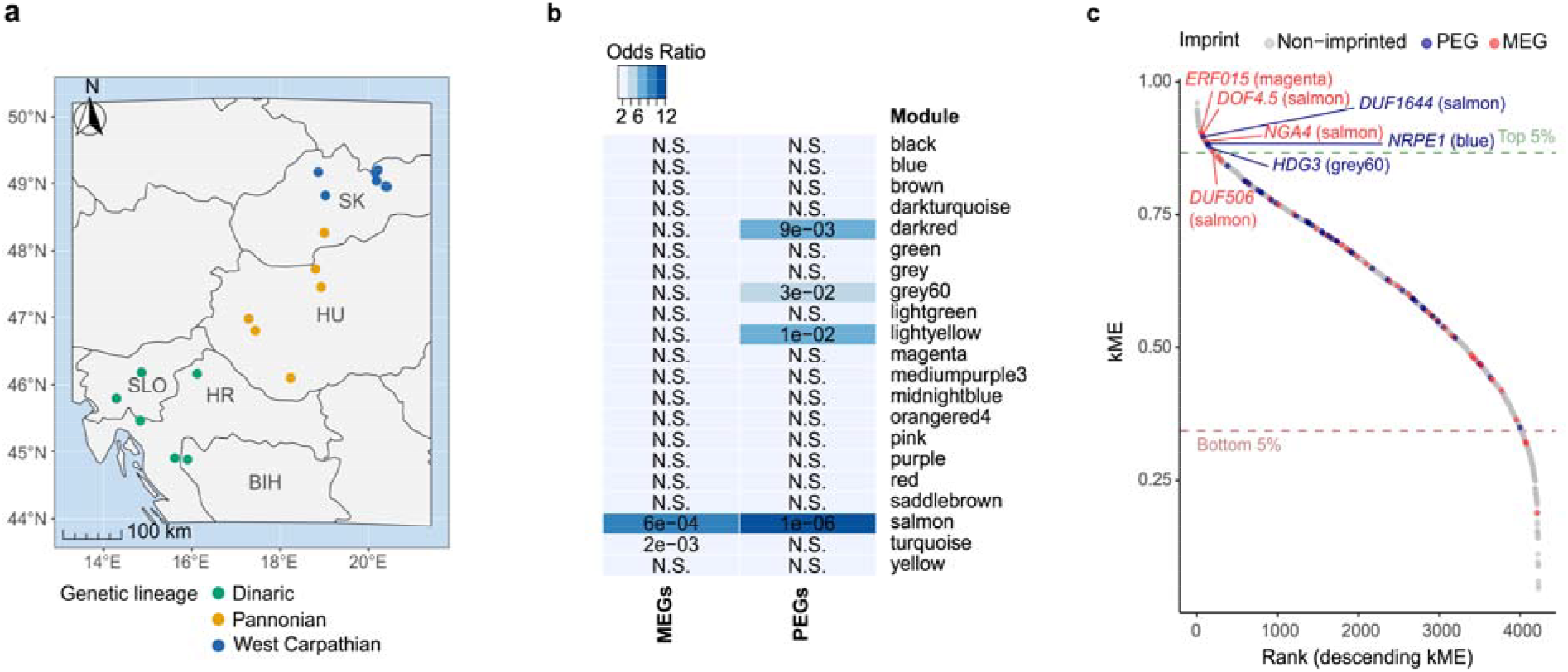
The species-level *A. arenosa* imprintome targets a handful of functional modules. (**a**) *A. arenosa* sampled populations, genetic lineages defined previously^10^. (**b**) Co-expression modules defined by WGCNA containing at least one imprinted gene (21 out of 45 modules). Odds ratio and overlap between a module’s genes and MEGs and PEGs calculated with GeneOverlap. N.S. denotes p-values > 0.05. (**c**) Ranked kME of genes upregulated in the endosperm compared to seed coat. Imprinted genes which placed in the top 5% of kME distribution are denoted by their functional name and module.

WGCNA builds a correlative network between genes and then splits it into modules, i.e., groups of genes with similar expression profiles, which are expected to be functionally related^11^. Our network had 45 functional modules, with five showing a significant enrichment in imprinted genes—hereafter imprinted modules (Fig. 1b). It is notable that MEGs and PEGs significantly co-occurred in one of the modules. This module, tagged as ‘salmon’, had the highest occurrence of both MEGs (five genes, p = 6.10^-4^) and PEGs (eight genes, p = 1.10^-6^), and was mostly enriched for biological process related to protein ubiquitination (Supplementary Fig. 1a). Imprinted genes encoding for components of the ubiquitin-26S proteasome pathway have been found previously^12^. The ubiquitin pathway has been identified to play an important role in the regulation of seed size^13,14^, which is largely determined by endosperm size^15^.

There were also parent-specific modules, indicating a degree of functional separation between PEGs and MEGs, in line with the different spatial occurrence of PEGs and MEGs in the endosperm^16^. The ‘turquoise’ was specifically enriched for MEGs (p = 0.002), while PEGs were significantly concentrated in the darkred (p = 0.009), grey60 (p = 0.03), and lightyellow (p = 0.01) modules (Supplementary Fig. 1b). The MEG-enriched ‘turquoise’ module was associated with biological processes related to cell division and cell cycle while the PEG-enriched ‘grey60’ module mirrored was enriched for DNA replication, DNA metabolic processes, and DNA replication initiation (Supplementary Fig. 1c). Cell cycle and DNA replication being tightly linked, this suggests a separated maternal and paternal influence over a crucial process for endosperm development. The PEG-enriched ‘darkred’ module showed a functional profile associated with cell killing and the disruption of cells of other organisms (Supplementary Fig. 1d). These functions most plausibly reflect receptor-like communication between tissues mediated by short signaling peptides with increasing evidence for their role in plant reproduction^17,18^. Finally, no GO terms met our statistical thresholds for the PEG-enriched ‘lightyellow’ module.

Overall, the clustering of imprinted genes to specific modules may pinpoint at pathways that are particularly dosage sensitive, and thus selection might act on the maintenance/rise of genomic imprinting in these pathways, rather than targeting specific genes. This would be consistent with the imprintome being only loosely conserved between species^19^, as establishing imprinting in a pathway would be more crucial than in a particular gene. Under a parental conflict scenario, these pathways may be the molecular arena for paternal and maternal influence to collide.

The connectivity of imprinted genes in a regulatory network can provide further insights into their role. Genes with low connectivity and pleiotropy are not as functionally constrained, while highly connected hub genes are subject to negative selection as any functional modifications can have severe consequences^20^. The kinship model assumes an ongoing arms race between maternal and paternal genomes. Genes subject to such an arms race are expected to usually have low connectivity and lie on the periphery of networks where there is more leeway for positive selection to act^21^. On the other hand, the sexual-key lock model expects imprinted genes to be strictly necessary for normal endosperm development and thus predicts they will be highly connected hub genes with pleiotropic effects.

To disentangle these two scenarios, we calculated the intramodular connectivity (kME) of imprinted genes and compared it with non-imprinted, endosperm-enriched, genes within their assigned modules. Neither MEGs nor PEGs as a group showed a significant difference in their intramodular connectivity compared to each other or to their non-imprinted counterparts (p = 0.48; Supplementary Fig. 2). As different imprinted genes may respond to different evolutionary pressures, we studied them individually, ranking all endosperm-upregulated genes by their absolute kME and identifying imprinted genes that placed within the top 5% of the connectivity distribution (Fig. 1c, Supplementary Table 3). Seven imprinted genes showed up as top 5% high-connectivity hubs, four MEGs and three PEGs. Notably, these hubs were distributed across several key functional modules, namely the ‘salmon’, ‘magenta’, ‘blue’, and ‘grey60’ modules. We propose that these imprinted hub genes act as keys and locks for endosperm development. Among them, *NRPE1*, a PEG encoding Pol V subunit, was found, highlighting an important role of the RdDM pathway in this key-lock system. Genes encoding a subunit of Pol IV and a common subunit of Pol IV & Pol V were found as PEGs in *A. thaliana* and *A. lyrata*, respectively^3,22^, suggesting the importance of the RdDM pathway being paternally imprinted across species. Among other imprinted gene hubs, *HDG3* was found as a PEG. This gene is also a PEG in *A. thaliana*, and variation in its imprinting was shown to affect seed weight and endosperm development^23^, suggesting a hub role too in *A. thaliana*. As a technical control, we compared the predicted targets of *HDG3* in our *A. arenosa* GRN with downregulated genes in *A. thaliana hdg3* mutant^23^ and found a significant overlap (hypergeometric test, p = 3.389982e-15, Supplementary Fig. 3).

Besides the role of imprinted genes in their network, the molecular interactions between parental genomes may shed light on how genomic imprinting contributes to endosperm development. These interactions might be direct, with imprinted genes regulating each other. To investigate this, we first identified 17 MEGs and 12 PEGs annotated as regulators of gene expression (including epigenetic, transcriptional, and post-transcriptional regulation; Supplementary Table 4). Using the R package GENIE3^24^, we determined the strength of regulatory links (edges) between these imprinted regulators and all the genes in the GRN. Targets of the 29 imprinted regulators were determined using the R package DIANE^25^ (Supplementary Table 5). We found that about one fourth of regulatory MEGs (4 out of 17) targeted more MEGs than expected by chance and two MEGs preferentially targeted PEGs (Fig. 2a). About one third of regulatory PEGs (4/12) preferentially targeted other PEGs, among which two PEGs also targeted more MEGs than expected by chance (Fig. 2b). Combining all regulatory edges together, paternal regulators were more than twice (2.24 odds ratio) as likely to target other PEGs than MEGs, relative to the number of regulatory opportunities for each (Fisher’s exact test, p = 0.012; Fig. 2c). These results echo a study on the paternally expressed MADS-box gene *PHE1* in *A. thaliana*, which was also found to target five other PEGs but only one MEG^9^. We show this trend is significant and applies more globally to regulatory PEGs. Maternal regulators, however, indiscriminately targeted both MEGs and PEGs (p = 0.8). The preference of PEG regulators for PEG targets was buffered by MEG regulators (more genes, indiscriminate targeting), reducing the consequences on the targets, which did not show any significant differences in the number of regulatory edges from PEG and MEG regulators (Fig. 2c). This molecular dynamics – the paternal genome driving its own impact on endosperm and the maternal genome buffering this influence – strikingly parallels the expectations of the parental conflict theory^2^.

**Fig. 2.**
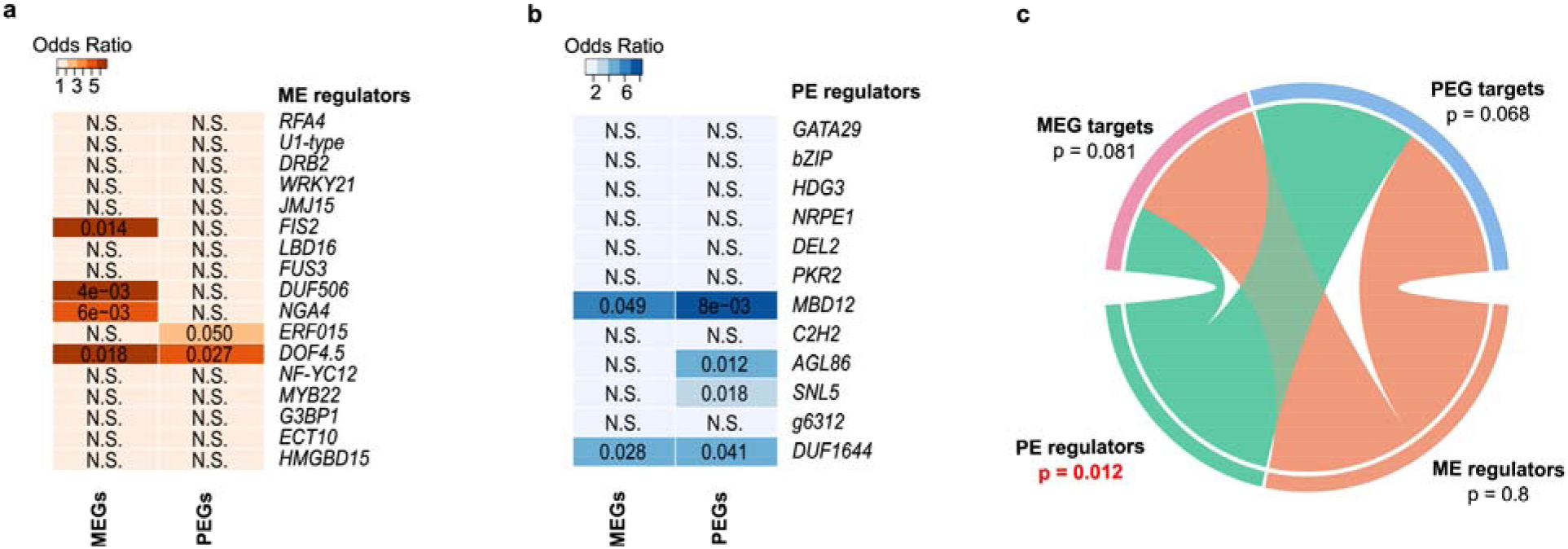
Direct interactions between imprinted genes show unilateral tendencies. (**a, b**) Odds ratio and overlap between the targets of imprinted regulators and MEGs/PEGs. (**a)** shows maternally expressed regulators and (**b**) shows paternally expressed regulators. Values in cells denote p-value, N.S. denotes p-values > 0.05 (Fisher’s exact test). (**c**) chord diagram representing edges (molecular relationship) between imprinted regulators and targets. The thickness of a chord is proportional to the number of edges. The p value (Fisher’s exact test) is relative to the comparison of within-vs between-parent interactions (e.g. proportion of PE regulator-PEG target edges vs PE regulator-MEG target edges, given the total number of potential edges for each category). ME: maternally expressed; PE: paternally expressed.

Another way MEGs and PEGs may interact is by regulating the same targets. To test this hypothesis, we analyzed shared targets among the regulatory imprinted genes. Because GENIE3 cannot distinguish indirect interactions (i.e. regulating a regulator of a target) from direct ones (Huynh-Thu et al., 2010; source), we disregarded target sharing where one regulator targeted the other one in the pair. With this criteria, significant target sharing happened in 14 pairs between regulatory MEGs and PEGs (Fig. 3a), while seven regulatory MEG-MEG pairs shared significantly more targets than expected by chance (Supplementary Fig. 4a), followed by PEG-PEG target sharing (five significant target overlaps; Supplementary Fig. 4b). This, however, merely reflected the opportunity for target sharing based on the number of regulators (17 maternal and 12 paternal regulators) and did not show a significant preference for either interparental or intraparental target sharing. Target sharing between imprinted regulators from different functional modules was rather common (Fig. 3b), suggesting a tight interconnection between pathways via the co-regulation of genes by PEGs and MEGs. For example, the MEG *JMJ15* and PEG *HDG3* significantly shared targets and belonged to the ‘turquoise’ (cell cycle) and ‘grey60’ (DNA replication) functional modules, respectively (Fig. 3b). This suggests a potential bridging between two related pathways brought by a paternal and maternal puzzle piece acting together. Given that *HDG3* is part of the top 5% hub genes and *JMJ15* also shows high connectivity (Fig. 1c; Supplementary Table 4), this pair could be a part of the sexual key-lock system essential to endosperm development.

**Fig. 3.**
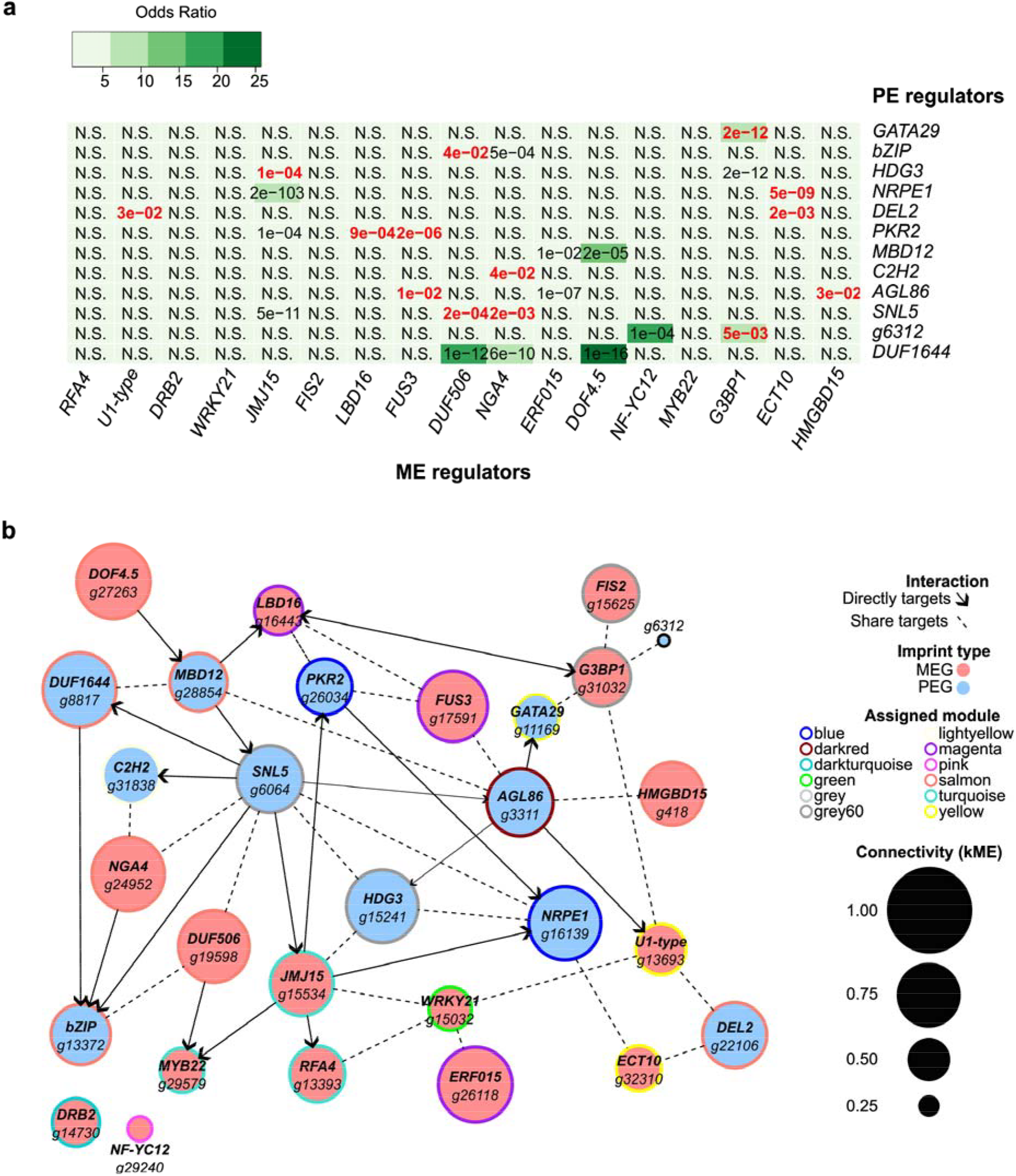
Paternal and maternal genes co-regulate targets. (**a**) Odds ratio and overlap between the targets of paternally and maternally expressed regulators calculated with GeneOverlap. N.S. denotes p-values > 0.05. P values in red show target sharing where none of the two regulators in the pair targets the other one, to avoid spurious inference of target sharing due to indirect regulation. (**b**) Schematic representation of the network of molecular relationships between PEGs and MEGs. When two genes shared targets and one regulated the other one, we represented the latter to avoid spurious inference of target sharing due to indirect regulation. When two genes shared targets and mutually targeted each other, we removed the interaction between the two genes as it is not possible to disentangle their relationship.

Taken together, we found that parental genomes are rather separated, even though several key imprinted regulators acted as interparental and intermodular bridges. This work provides a comprehensive picture of how parental conflict and collaboration happen at the molecular level and how both processes can coexist and shape endosperm development.

## Methods

### Plant material

Seeds of *Arabidopsis arenosa* were collected from populations in its natural range (Central and Southern Europe) between 2018 and 2022. The GPS locations are provided in Supplementary Table 1. Seeds were surface-sterilized using a solution containing 5% sodium hypochlorite and 0.01% (v/v) Triton X-100. After sterilization, the seeds were planted on Petri dishes with Murashige and Skoog medium (1/2× MS salts (DUCHEFA BIOCHEMIE B.V M0222), 0.8% plant agar (DUCHEFA BIOCHEMIE B.V P100), 10 mM MES (DUCHEFA BIOCHEMIE B.V M150), pH 5.8). Planted seeds were stratified for 7 days at 4 °C in the dark and then germinated under controlled conditions of 21/18 °C with a 16/8 hour day/night cycle for 2 weeks. After germination, seedlings were transferred to soil and cultivated in the same conditions for another 5 weeks. Subsequently, plants were vernalized for at least two months at 4 °C with an 8/16 hour day/night cycle. Vernalized plants were then grown under long-day conditions (21/18 °C with a 16/8 hours day/night cycle) to induce flowering.

### Endosperm isolation

To maximize the number of SNPs between parents, crosses were performed between plants of different populations (see Supplementary Table 1 for the details of the crossing design). Crosses were done by emasculating the unopened flower buds of the maternal plant and transferring pollen from the paternal plant 1-3 days after emasculation to the pistil. Siliques were collected 10-16 days after pollination and endosperm, embryo and seed coat were manually isolated and separated in the developmental stage, varying from early to late torpedo stage, all at room temperature (∼24 °C). Siliques were placed on 3M micropore tape, opened by tweezer (Precision tweezers DUMONT® straight with extra fine tips Dumoxel®, 5) and individual seeds were transferred into extraction buffer (0.3 M D(-)-Sorbitol (Millipore® 1.07758), 5 mM MES (DUCHEFA BIOCHEMIE B.V M1503), pH 5.7, filter sterilized). Using the tweezer and insulin syringe (Omnican® 100), individual parts of the seeds were separated. Endosperm and embryo samples originated from 120-160 seeds, and seed coat samples from 240-320 seeds. Individual fractions were transferred into 2.0 ml safe-lock tubes (Eppendorf 0030123344) and centrifuged for 90 sec at 12,000 g in the bench MiniSpin® centrifuge. Supernatant from all samples was discarded. Endosperm samples were frozen immediately in liquid nitrogen, and embryos and seed coats were washed two times in 1 ml of extraction buffer before freezing. All samples were kept at -80 °C for further use.

### RNA & DNA sequencing

RNA was isolated using the RNAqueous™ Total RNA Isolation Kit (Thermo Fisher Scientific AM1912) with the inclusion of Plant RNA Isolation Aid (Thermo Fisher Scientific AM9690) in the lysis buffer. DNA was extracted from leaves using DNeasy® Plant Mini Kit (Qiagen 69104).

Total RNA and DNA samples were sent to Novogene (UK) Company Limited for library preparation and sequencing. RNA samples undergo quality control, poly A enrichment, directional mRNA library preparation, and Illumina Sequencing PE150, with raw data obtained 4.5 G and 7.5 G. DNA samples undergo quality control, whole genome library preparation (350 bp), and Illumina Sequencing PE150, with raw data obtained 6 G.

### Gene annotation

Structural gene annotation of *Arabidopsis arenosa* genome assembly was done using Augustus version 3.3.226. The full-length coding sequences from *Arabidopsis lyrata* and training annotation files of *Arabidopsis thaliana* were used for homology-based gene prediction, resulting in 33,911 gene models. Functional annotation of predicted protein sequences was done using 21,867 and 23,926 reciprocal blast hits (e-val less than 0.001) from *Arabidopsis thaliana* and *Arabidopsis lyrata*, respectively (Supplementary Table 6).

### Variant calling and identification of imprinted genes

Paired-end 150 bp sequenced reads from both parental DNA and offspring RNA were mapped to the *A. arenosa* reference genome (GCA_026151155.1) using BWA version 0.7.17^27^ with a smart pairing option for paired-end alignment. Duplicates were then removed and SNPs called using GATK version 4.2.5.0^28^. The obtained SNPs were filtered by genome coverage (DP > 7) and genotype quality (GQ > 19). SNPs within exons and without a common parental allele were further used for the identification of imprinted genes. Each SNP was tested for significant (p-adj < 0.05) deviation from the expected ratio of maternal to paternal allele 2:1 in both reciprocal crosses using the Binomial test. Imprinted genes were identified when a gene contained at least two SNPs showing significant parent-of-origin bias in the same direction. Additionally, we took into consideration that at least two-thirds of all informative SNPs within the gene (regardless of significance) exhibited the same parental bias.

Finally, we removed genes that might represent contamination from maternal tissue rather than imprinted genes. To do so, the raw reads from endosperm and seed coat samples were trimmed for adaptors using Trim Galore version 0.4.1^29^ and aligned to the *A. arenosa* reference genome (GCA_026151155.1) using HiSat2 version 2.1.0^30^. Reads were then assigned to features and meta-features using Subread version 1.5.231 according to the prepared genome annotation. Differential expression analysis was performed using DESeq2 version 1.38.3^32^ in R v.4.2.2 to compare gene expression between endosperm and seed coat samples. Genes identified as parentally biased and significantly upregulated in endosperm compared to seed coat (Benjamini–Hochberg FDR-adjusted P-value < 0.05) were kept as imprinted genes.

### WGCNA network construction

All the RNA-Seq samples were re-processed for the gene regulatory network construction. The raw reads were filtered using FASTX-Toolkit v3.3^33^ and Trimmomatic v0.39^34^ to remove the unstable nucleotide bases from the beginning of the sequence and the adapters, respectively. Before proceeding to mapping, the quality of trimmed reads was rechecked to confirm the efficiency of the trimming step. Subsequently, reads were mapped to the *Arabidopsis arenosa* reference genome (GCA_026151155.1) using STAR v2.7^35^ with adjusted parameter for --outFilterMultimapNmax: 1. This option was used to retain the only uniquely mapped reads for further analysis. Finally, uniquely aligned reads with features were counted for each gene using HTSeq v0.11.1^36^. Read counting was performed for all reads with default parameters except -m intersection-nonempty -s no -r pos. *Arabidopsis arenosa* genome annotation generated in the “gene annotation” section was used for counting uniquely mapped reads.

We then built three weighted gene correlative networks using the “WGCNA” package^11^ in R version 4.3.1 based on the transcriptomes from the same 62 endosperm samples used to identify imprinted genes. In brief, WGCNA constructs a global network based on correlation, which is then divided into clusters of highly interconnected genes termed modules. Genes within a module are expected to be functionally related. Prior to network construction, the transcriptome data were TPM normalized, filtered for genes with high counts (expressed >0.5 TPM in at least 3 samples), and log-transformed for variance stabilization [log_2_(x + 1)]. Prior to analysis, the data were visually checked for outliers in a hierarchical clustering dendrogram and PCA. No outliers were found; hence, we moved forward with all our data. We used the soft thresholding power of 14 (the power to which co-expression similarities are raised to enhance strong and diminish weak correlation) in all three networks. In each case, we chose a thresholding power that exceeded a scale-free topology fit index of 0.8 and had a small mean connectivity (<200). The signed networks were constructed using the function blockwiseModules() with a maximum block size of 28,000 genes. We used biweight mid-correlation “bicor,” a robust alternative to Pearson correlation, and set maxPOutliers to 0.05 to minimize the effects of outliers. The TOMType argument was set to “signed.” The dendrogram cut height for the merging of similar modules, mergeCutHeight, was set to 0.25.

### Characterizing modules with imprinted genes

Modules with a significant number of imprinted genes, hereafter ‘imprinted’ modules, were identified using the package “GeneOverlap” in R version 4.3.1, which uses a Fisher’s exact test to find significant overlaps between module genes and imprinted genes^37^. We performed GO term enrichment on the *Arabidopsis thaliana* homologs of the imprinted modules’ *A. arenosa* genes with a custom background of *A. thaliana* homologs of all *A. arenosa* genes using PlantRegMap, a plant regulatory data and analysis platform^38^. We enriched for biological process (BP), molecular function (MF), and cellular component (CC) with a threshold p-value ≤ 0.05.

### Gene connectivity

The connectivity of genes within their modules was determined using the WGCNA R package function signedKME(), which calculates the correlation between a gene’s expression pattern and a module’s eigengene, the first principle component of a module’s expression profile. This correlation is known as intramodular connectivity (kME). The higher a gene’s kME in a module, the more connected we can expect it to be in the given module. To compare connectivity between groups of genes, we used the absolute value of a gene’s kME within its assigned module. We compared the kME of MEGs, PEGs, and all non-imprinted genes belonging to modules with at least one imprinted gene using the Kruskal-Wallis test followed by a post-hoc Tukey-Kramer test.

### Inference of regulatory links between genes

We used the package “GENIE3” in R version 4.3.1 with default parameters to calculate the strength of regulatory links between genes^24^. We selected only genes with genes upregulated in the endosperm compared to the seed coat, leaving us with 4820 genes. From these genes, we identified *A. arenosa* genes with *A. thaliana* homologs involved in gene expression regulation (including transcription, translation, and epigenetic regulation) using The Arabidopsis Information Resource (TAIR) GO Annotations. From these genes, 29 were imprinted and chosen as regulators for our GENIE3 analysis (17 MEGs, 12 PEGs; Supplementary Table 4). We used a weighted threshold calculated by the function test_edges() from the R package “DIANE” to predict regulator target genes^25^. The global network density was chosen as 0.1, and an FDR correction of 0.05 was applied. To see whether MEGs and PEGs share targets among and between each other, we used the R package “GeneOverlap.”

## Code availability

The customized codes used in this study are available from the corresponding author upon request.

## Acknowledgements

The authors appreciate F. Kolář for giving *A. arenosa* seeds. The authors are grateful to Daniel Lang for his invaluable feedback on the design and analyses along the project. This work was funded by the Czech Science Foundation (22-20240S to CLP). MK and AlPe were partly funded by the MEYS INTER-COST project LUC24011. Computational resources were provided by the e-INFRA CZ project (ID:90254), supported by the Ministry of Education, Youth and Sports of the Czech Republic.

## Author contributions

TB and CLP designed the research; VČ and AdPř performed the experiments; TB, MK, SS and ÖI analyzed the data; TB and CLP wrote the paper; all authors reviewed and modified the manuscript.

## References

1. Batista, R. A. & Köhler, C. Genomic imprinting in plants-revisiting existing models. Genes Dev 34, 24–36 (2020).

2. Haig, D. & Westoby, M. Parent-Specific Gene-Expression and the Triploid Endosperm. American Naturalist 10.1086/284971 (1989) doi:10.1086/284971.

3. Gehring, M., Missirian, V. & Henikoff, S. Genomic Analysis of Parent-of-Origin Allelic Expression in Arabidopsis thaliana Seeds. PLOS ONE 6, e23687 (2011).

4. Figueiredo, D. D., Batista, R. A., Roszak, P. J. & Köhler, C. Auxin production couples endosperm development to fertilization. Nature Plants 1, 1–6 (2015).

5. Chaudhury, A. M. et al. Fertilization-independent seed development in Arabidopsis□thaliana. Proceedings of the National Academy of Sciences 94, 4223–4228 (1997).

6. Figueiredo, D. D. & Köhler, C. Auxin: a molecular trigger of seed development. Genes Dev 32, 479–490 (2018).

7. Grossniklaus, U., Vielle-Calzada, J.-P., Hoeppner, M. A. & Gagliano, W. B. Maternal Control of Embryogenesis by MEDEA, a Polycomb Group Gene in Arabidopsis. Science 280, 446–450 (1998).

8. Wolff, P., Jiang, H., Wang, G., Santos-González, J. & Köhler, C. Paternally expressed imprinted genes establish postzygotic hybridization barriers in Arabidopsis thaliana. eLife 4, e10074 (2015).

9. Batista, R. A. et al. The MADS-box transcription factor PHERES1 controls imprinting in the endosperm by binding to domesticated transposons. eLife 8, e50541.

10. Kolář, F. et al. Northern glacial refugia and altitudinal niche divergence shape genome-wide differentiation in the emerging plant model Arabidopsis arenosa. Molecular Ecology 25, 3929–3949 (2016).

11. Langfelder, P. & Horvath, S. WGCNA: an R package for weighted correlation network analysis. BMC Bioinformatics 9, 559 (2008).

12. Hsieh, T.-F. et al. Regulation of imprinted gene expression in Arabidopsis endosperm. Proceedings of the National Academy of Sciences 108, 1755–1762 (2011).

13. Xia, T. et al. The Ubiquitin Receptor DA1 Interacts with the E3 Ubiquitin Ligase DA2 to Regulate Seed and Organ Size in Arabidopsis. Plant Cell 25, 3347–3359 (2013).

14. Du, L. et al. The Ubiquitin Receptor DA1 Regulates Seed and Organ Size by Modulating the Stability of the Ubiquitin-Specific Protease UBP15/SOD2 in Arabidopsis. Plant Cell 26, 665–677 (2014).

15. Garcia, D., Fitz Gerald, J. N. & Berger, F. Maternal Control of Integument Cell Elongation and Zygotic Control of Endosperm Growth Are Coordinated to Determine Seed Size in Arabidopsis. Plant Cell 17, 52–60 (2005).

16. Picard, C. L., Povilus, R. A., Williams, B. P. & Gehring, M. Transcriptional and imprinting complexity in Arabidopsis seeds at single-nucleus resolution. Nat. Plants 7, 730–738 (2021).

17. Zhu, S., Fu, Q., Xu, F., Zheng, H. & Yu, F. New paradigms in cell adaptation: decades of discoveries on the CrRLK1L receptor kinase signalling network. New Phytologist 232, 1168–1183 (2021).

18. İltaş, Ö. et al. The abundance of pollen coat small signaling proteins shows limited convergence between independent selfing transitions in Arabidopsis and Capsella. New Phytologist 250, 3460–3474 (2026).

19. Hatorangan, M. R., Laenen, B., Steige, K. A., Slotte, T. & Köhler, C. Rapid Evolution of Genomic Imprinting in Two Species of the Brassicaceae. Plant Cell 28, 1815–1827 (2016).

20. Josephs, E. B., Wright, S. I., Stinchcombe, J. R. & Schoen, D. J. The Relationship between Selection, Network Connectivity, and Regulatory Variation within a Population of Capsella grandiflora. Genome Biol Evol 9, 1099–1109 (2017).

21. Molodtsova, D., Harpur, B. A., Kent, C. F., Seevananthan, K. & Zayed, A. Pleiotropy constrains the evolution of protein but not regulatory sequences in a transcription regulatory network influencing complex social behaviors. Front. Genet. 5, (2014).

22. Klosinska, M., Picard, C. L. & Gehring, M. Conserved imprinting associated with unique epigenetic signatures in the Arabidopsis genus. Nat Plants 2, 16145 (2016).

23. Pignatta, D., Novitzky, K., Satyaki, P. R. V. & Gehring, M. A variably imprinted epiallele impacts seed development. PLOS Genetics 14, e1007469 (2018).

24. Huynh-Thu, V. A., Irrthum, A., Wehenkel, L. & Geurts, P. Inferring Regulatory Networks from Expression Data Using Tree-Based Methods. PLOS ONE 5, e12776 (2010).

25. Cassan, O., Lèbre, S. & Martin, A. Inferring and analyzing gene regulatory networks from multi-factorial expression data: a complete and interactive suite. BMC Genomics 22, 387 (2021).

26. Stanke, M., Diekhans, M., Baertsch, R. & Haussler, D. Using native and syntenically mapped cDNA alignments to improve de novo gene finding. Bioinformatics 24, 637–644 (2008).

27. Li, H. Aligning sequence reads, clone sequences and assembly contigs with BWA-MEM. arXiv:1303.3997 [q-bio] http://arxiv.org/abs/1303.3997 (2013).

28. McKenna, A. et al. The Genome Analysis Toolkit: a MapReduce framework for analyzing next-generation DNA sequencing data. Genome Res 20, 1297–1303 (2010).

29. Martin, M. Cutadapt removes adapter sequences from high-throughput sequencing reads. EMBnet.journal 17, 10–12 (2011).

30. Kim, D., Paggi, J. M., Park, C., Bennett, C. & Salzberg, S. L. Graph-based genome alignment and genotyping with HISAT2 and HISAT-genotype. Nat Biotechnol 37, 907–915 (2019).

31. Liao, Y., Smyth, G. K. & Shi, W. The Subread aligner: fast, accurate and scalable read mapping by seed-and-vote. Nucleic Acids Res 41, e108 (2013).

32. Love, M. I., Huber, W. & Anders, S. Moderated estimation of fold change and dispersion for RNA-seq data with DESeq2. Genome Biol 15, 550 (2014).

33. Gordon, A. & Hannon, G. FASTX-Toolkit. FASTQ/A short-reads pre-processing tools. (2010).

34. Bolger, A. M., Lohse, M. & Usadel, B. Trimmomatic: a flexible trimmer for Illumina sequence data. Bioinformatics 30, 2114–2120 (2014).

35. Dobin, A. et al. STAR: ultrafast universal RNA-seq aligner. Bioinformatics 29, 15–21 (2013).

36. Anders, S., Pyl, P. T. & Huber, W. HTSeq—a Python framework to work with high-throughput sequencing data. Bioinformatics 31, 166–169 (2015).

37. Shen, L. & Sinai IsoMaM. GeneOverlap: Test and visualize gene overlaps. Bioconductor 10.18129/B9.bioc.GeneOverlap (2026).

38. Tian, F., Yang, D.-C., Meng, Y.-Q., Jin, J. & Gao, G. PlantRegMap: charting functional regulatory maps in plants. Nucleic Acids Res 48, D1104–D1113 (2020).

