## Supplementary Fig for "Imprinted regulatory networks reveal the molecular crosstalk between paternal and maternal genomes in the endosperm of *Arabidopsis arenosa*"

**Supplementary Figures**

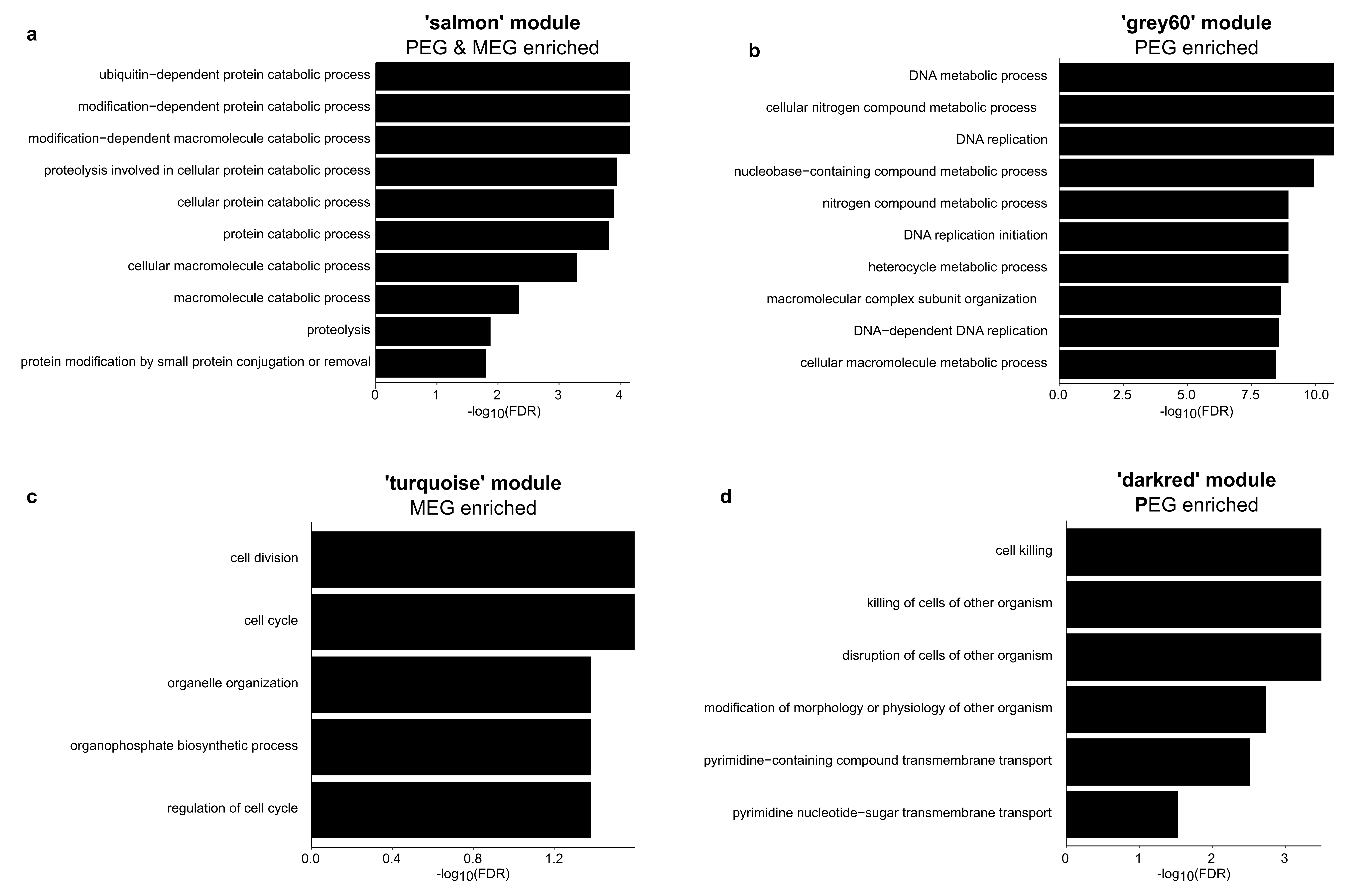

**Supplementary Figure 1. Genomic imprinting affects a handful of functional modules.** Enriched Biological processes GO terms for the (**a**) ‘salmon’ module, which is enriched for both MEGs and PEGs, (**b**) ‘grey60’ module, which is enriched for PEGs, (**c**) ‘turquoise’ module, which is enriched for MEGs and (**d**) ‘darkred’ module, which is enriched for PEGs.

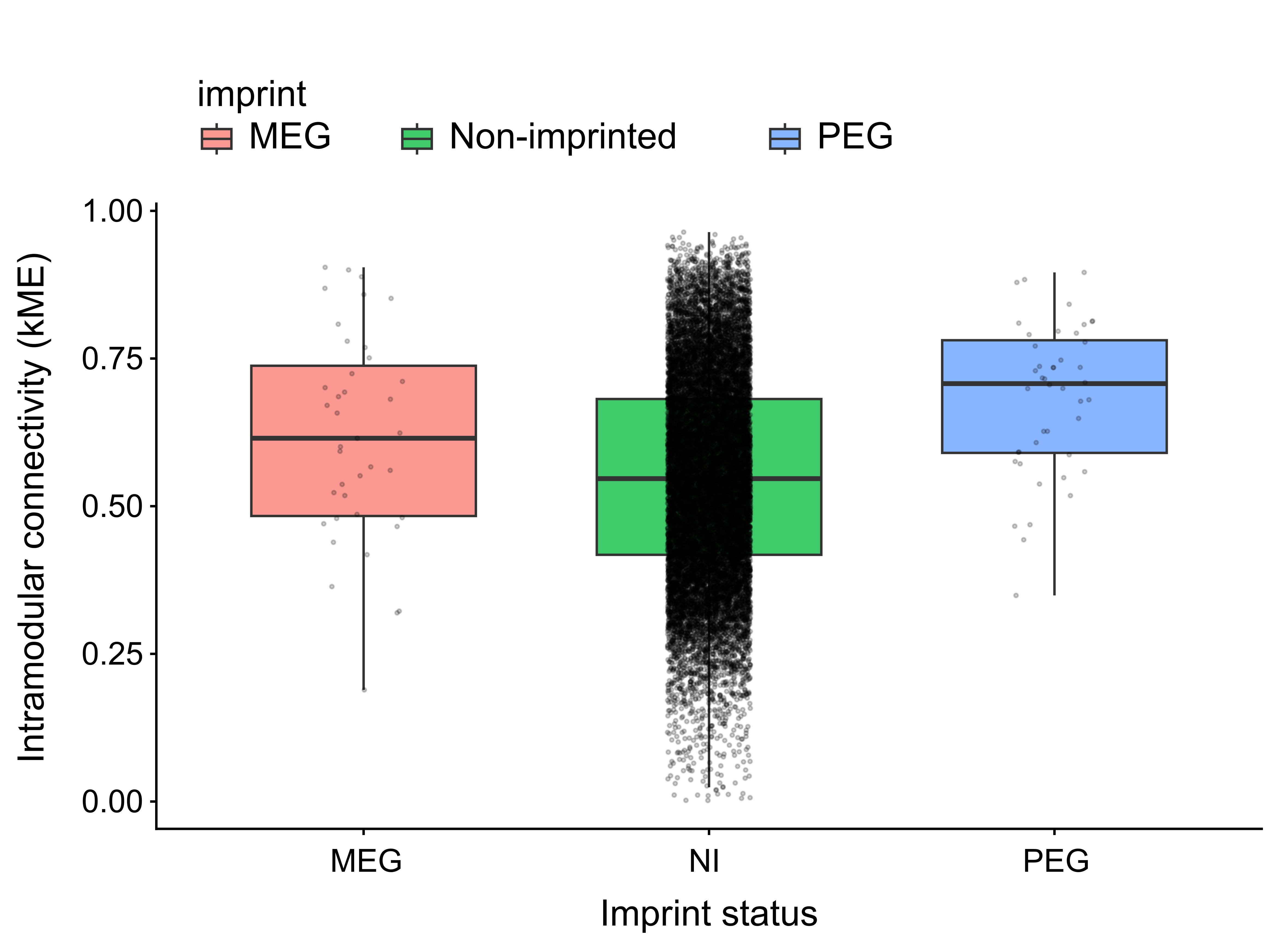

**Supplementary Figure 2. Imprinted genes as a group do not act as hubs.** Comparison of intramodular connectivity between MEGs, PEGs, and non-imprinted genes (NI). Non-imprinted genes only include genes upregulated in the endosperm compared to the seed coat (n = 4820 genes). No significant differences between gene groups were found (Fisher’s exact test, p > 0.05).

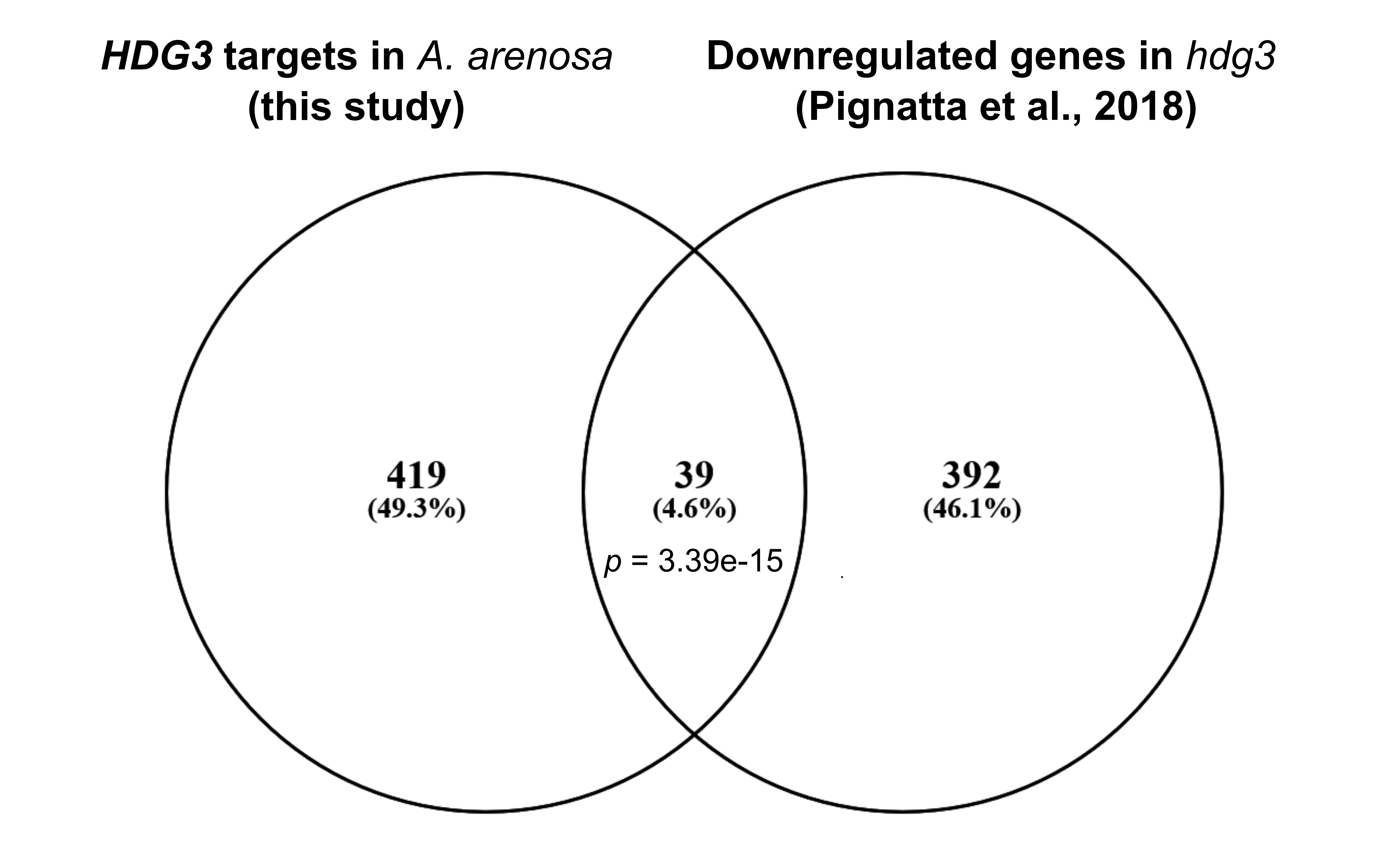

**Supplementary Figure 3. Gene network predictions and experimental data coincide.** Comparison of the genes predicted as targets of *HDG3* in *A. arenosa* and the genes downregulated in the hdg3 mutant in *A. thaliana*^8^. The p value refers to a hypergeometric test. *A. thaliana* gene IDs were used to make this comparison.

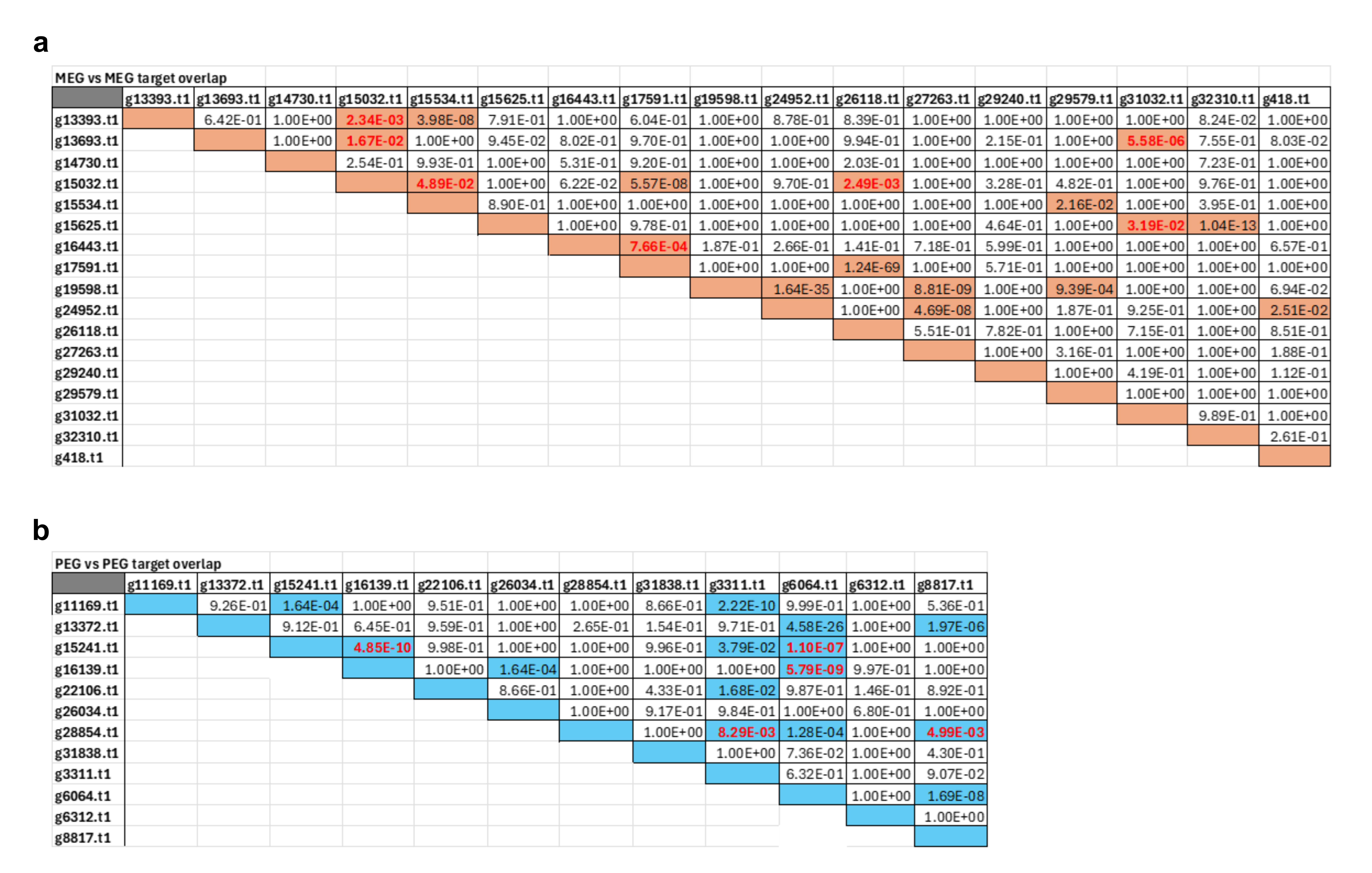

**Supplementary Figure 4. Paternal and maternal genes co-regulate targets.** (**a**) Odds ratio and overlap between the targets of maternally expressed regulators calculated with GeneOverlap. (**b**) Odds ratio and overlap between the targets of paternally expressed regulators calculated with GeneOverlap. N.S. denotes p-values > 0.05 (Fisher’s exact test). P values in red show target sharing where none of the two regulators in the pair targets the other one, to avoid spurious inference of target sharing due to indirect regulation.
